# Chromosome-level genome of *Poropuntius huangchuchieni* provides a diploid progenitor-like reference genome for the allotetraploid *Cyprinus carpio*

**DOI:** 10.1101/2020.09.25.312652

**Authors:** Lin Chen, Bijun Li, Baohua Chen, Chengyu Li, Zhixiong Zhou, Tao Zhou, Weidi Yang, Peng Xu

## Abstract

The diploid *Poropuntius huangchuchieni* in the cyprinid family, which is distributed in the Mekong and Red River basins, is one of the most closely related diploid progenitor-like species of allotetraploid common carp. Therefore, the *P. huangchuchieni* genome is essential for polyploidy evolution studies in Cyprinidae. Here, we report a chromosome-level genome assembly of *P. huangchuchieni* by integrating Oxford Nanopore and Hi-C technology. The assembled genome size was 1021.38 Mb, 895.66 Mb of which was anchored onto 25 chromosomes with a N50 of 32.93 Mb. The genome contained 486.28 Mb repetitive elements and 24,099 protein-coding genes. Approximately 95.9% of the complete BUSCOs were detected, suggesting a high completeness of the genome. Evolutionary analysis revealed that *P. huangchuchieni* diverged from *Cyprinus carpio* at approximately 12 Mya. Genome comparison between *P. huangchuchieni* and the B subgenome of *C. carpio* provided insights into chromosomal rearrangements during the allotetraploid speciation. With the complete gene set, 17,474 orthologous genes were identified between *P. huangchuchieni* and *C. carpio*, providing a broad view of the gene component in the allotetraploid genome, which is critical for future genomic and genetic analyses. The high-quality genomic dataset created for *P. huangchuchieni* provides a diploid progenitor-like reference for the evolution and adaptation of allotetraploid carps.

## INTRODUCTION

*Poropuntius huangchuchieni* is a medium-sized economic freshwater fish species, belonging to the *Poropuntius* genus, which has a broad geographic distribution spanning from southwestern China to Myanmar, Thailand, the Malay Peninsula, and Sumatra (Kottelat, 2013) with the Mekong River System being the most likely center of diversity for the genus. In China, the genus is naturally distributed in the Yunnan Plateau, the southeastern neighbor of the Qinghai-Tibetan Plateau, which consistently responded to the Qinghai-Tibetan Plateau’s (QTP) uplifting during the Pliocene epoch, resulting in a dramatic change in the region’s climate (Cheng, Liu, Gao, Tang, & Yue, 2001; Ming & Shi, 2006). Consequently, the aquatic species in these regions may have experienced a driving force for diversification and speciation in evolutionary history, and their genetic structure and distributions may, in turn, provide a natural link in understanding the concurrent geographical and biotic evolution of a given region (Dubut et al., 2012). It has been reported that the phylogenetic pattern of *P. huangchuchieni* is mostly associated with the drainage structures and geomorphological history of the Southwest Yunnan Plateau (Wu et al., 2013). Additionally, the differentiation of the major evolutionary lineages among the Mekong, Salween, and Red River systems coincides with the Kunlun-Yellow River Movement (Wu et al., 2013). However, the lack of whole genome data has limited further interests on genetic and evolutionary studies.

In particular, as a member of the Barbinae subfamily, Cyprinid family, the Poropuntius is one of the most closely related diploid progenitor-like species to common carp *(Cyprinus carpio*), which has an allotetraploidized genome that was generated by merging of two diploid genomes during evolution (Xu et al., 2019), thereby making it a valuable resource for genetic and evolutionary studies of polyploid origins and adaptations in Cyprinidae. Cyprinids is a diverse teleost family with more than 2,400 species (Winfield & Nelson, 2012), of which the subfamily Cyprininae comprises over 1,300 species with karyotypes ranging from 2n=50 to ~470, including barbine, cyprinine, labeonine and schizothoracine (Arai, 2011). To date, several polyploidy genomes in Cyprinidae have been developed, including the three *Sinocyclocheilus* cavefish (Yang et al., 2016), goldfish *(Carassius auratus)* (Chen et al., 2019; Luo et al., 2020), *Oxygymnocypris stewartii* (Liu et al., 2019) and the common carp (Xu et al., 2019; Xu et al., 2014). Unlike polyploidy plants, such as the octoploid strawberry (Edger et al., 2019) and the allopolyploid *Brassica napus* (Lu et al., 2019), the origin and polyploid evolution of allotetraploid carps have not been elucidated. Evolutionary analysis of polyploid animals is more challenging because of the lack of progenitor-like genome data. The polyploidy of the cyprinid species is an ideal model for ploidy evolution studies of teleost fish. The allotetraploid *C. carpio* genome was divided into A and B subgenomes based on the genome identity between *P. huangchuchieni* and *C. carpio*, and the B subgenome of *C. carpio* was found to be derived from the Barbinae approximately 12.4 Mya (Xu et al., 2019). The successful identification of the progenitor-like lineageof *C. carpio* highlighted the importance of chromosome-level genome data for polyploid evolution studies in Cyprinidae.

In this study, we assembled and annotated a chromosome level reference genome for *P. huangchuchieni*, and conducted genome comparisons between *P. huangchuchieni* and *C. carpio*. We aimed to provide a foundation for studies of polyploid origin, speciation and adaptation in polyploidy Cyprinidae.

## Results and Discussion

### Estimation of genome size

Illumina short reads libraries from *P. huangchuchieni* (Figure 1) were constructed and sequenced on the Hiseq X ten platform, which produced a total of 57.35 Gb of raw data. After quality filtering, 49.14 Gb clean data was obtained for genome size estimation (Table 1, Table S1). Based on a 17-kmer analysis and a dominant peak depth of 47, the genome size was estimated to be 934.05 Mb, which was similar to that in other diploid species in Cyprinidae, such as *Onychostoma macrolepis* and grass carp *(Ctenopharyngodon idellus)* (Sun et al., 2020; Wang et al., 2015). The heterozygosity was estimated to be 0.68% (Table S2).

**Table 1.** Sequencing data of *P. huangchuchieni* for genome assembly and annotation.

| Sequencing technology | Raw data (Gb) | Clean data (Gb) | Mean read length (bp) |
| --- | --- | --- | --- |
| Hiseq survey | 57.35 | 49.14 | 150 |
| Nanopore | 76.06 | 74.78 | 23,641 |
| Hi-C | 100.39 | 99.28 | 150 |
| RNA-Seq | 89.98 | 88.41 | 150 |

**Figure 1.**
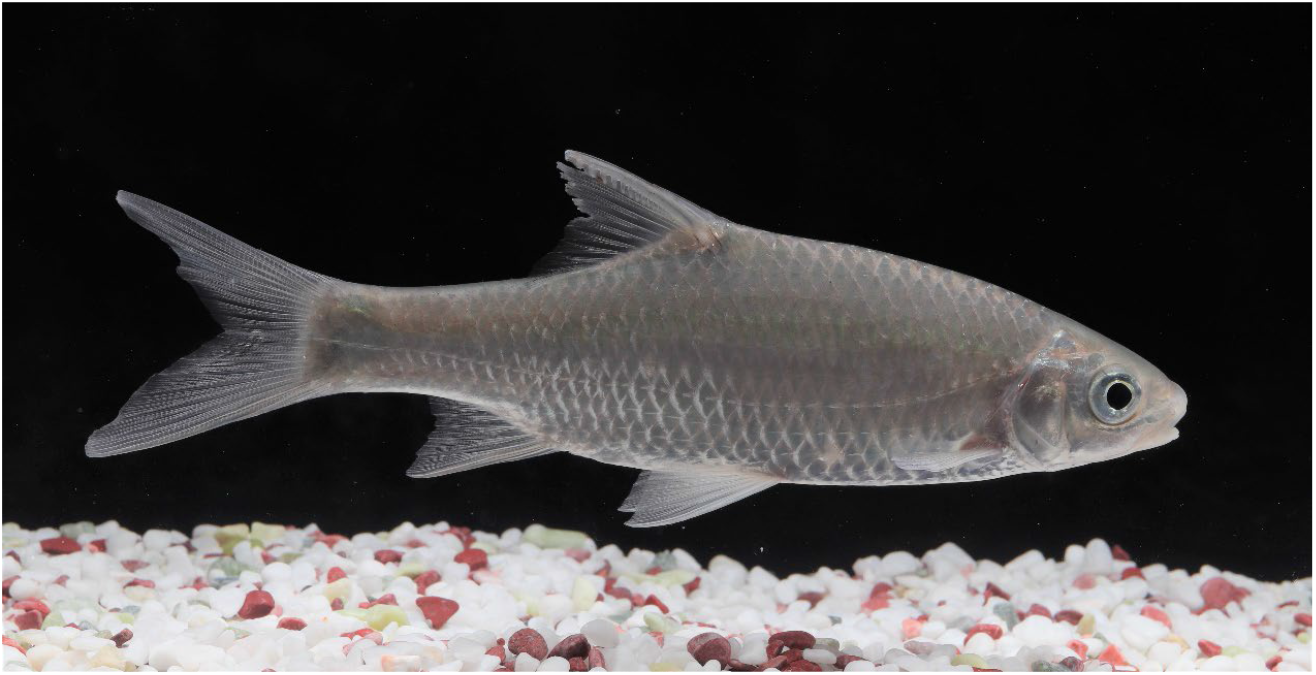
A photograph of *P. huangchuchieni*.

### *De novo* assembly of the *P. huangchuchieni* reference genome

Single-molecule sequencing with Nanopore technology generated 76.06 Gb raw reads, and 74.78 Gb clean data were retained after quality control (Table 1). The average read length and N50 of read length were 21.04 kb and 30.17 kb, respectively (Table S3). The advantage of using Nanopore technology with long consecutive reads contributed to the high quantity genome assembly of *P. huangchuchieni* (Table S4).

The Nanopore long consecutive reads were assembled *de novo* and were corrected by mapping the Illumina short reads to the contigs. A preliminary genome assembly with 919 contigs and an N50 of 4.10 Mb was generated (Table S5). The genome size was approximately 1020.36 Mb, which was close to the estimated genome size.

### Chromosomal-level assembly and synthetic analysis

A total of 100.39 Gb Hi-C raw reads was obtained, and 99.28 Gb was retained after quality control (Table S6). The contigs in the Nanopore draft assembly were then anchored and oriented into a chromosomal-scale assembly using the Hi-C scaffolding approach. Ultimately, we obtained a draft genome assembly of 1021.38 Mb in length, with a scaffold N50 value of 32.93 Mb (Figure 2A and Table 2). The genome assembly contained 25 chromosomes, with chromosome lengths ranging from 26.38 Mb to 74.78 Mb, and covered 895.66 Mb (87.69%) of the *P. huangchuchieni* assembly (Table S7). To further evaluate the completeness of the genome assembly, we checked the gene content with the BUSCO software using 4,584 single-copy genes conserved in Actinopterygii. BUSCO analysis showed that the assembly retrieved 95.9% (4,399/4,584) of the conserved single-copy ortholog genes (Table 3). This evidence supports the high-quality assembly of the *P. huangchuchieni* genome.

**Table 2.**
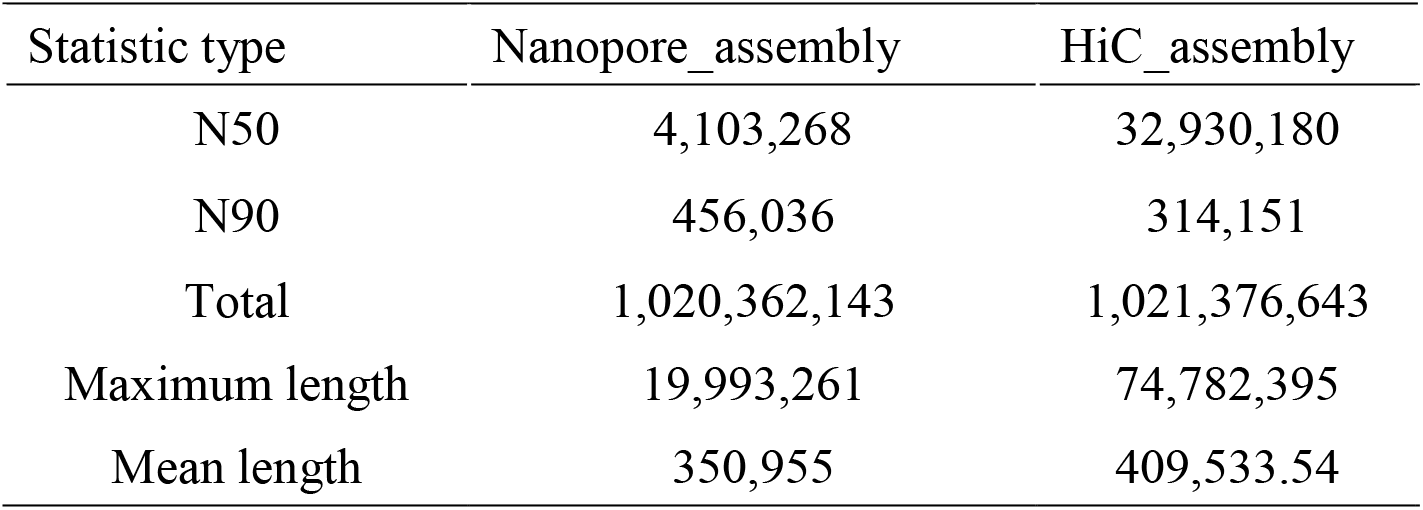
Statistics of the *P. huangchuchieni* genome assembly.

| Statistic type | Nanopore_assembly | HiC_assembly |
| --- | --- | --- |
| N50 | 4,103,268 | 32,930,180 |
| N90 | 456,036 | 314,151 |
| Total | 1,020,362,143 | 1,021,376,643 |
| Maximum length | 19,993,261 | 74,782,395 |
| Mean length | 350,955 | 409,533.54 |

**Table 3.** Assessment of genome completeness by BUSCO.

| Type | Number | Percentage (%) |
| --- | --- | --- |
| Complete BUSCOs (C) | 4,399 | 95.9 |
| Complete and single-copy BUSCOs (S) | 4,201 | 91.6 |
| Complete and duplicated BUSCOs (D) | 198 | 4.3 |
| Fragmented BUSCOs (F) | 64 | 1.4 |
| Missing BUSCOs (M) | 121 | 2.7 |
| Total BUSCO groups searched | 4,584 | 100 |

**Figure 2.**
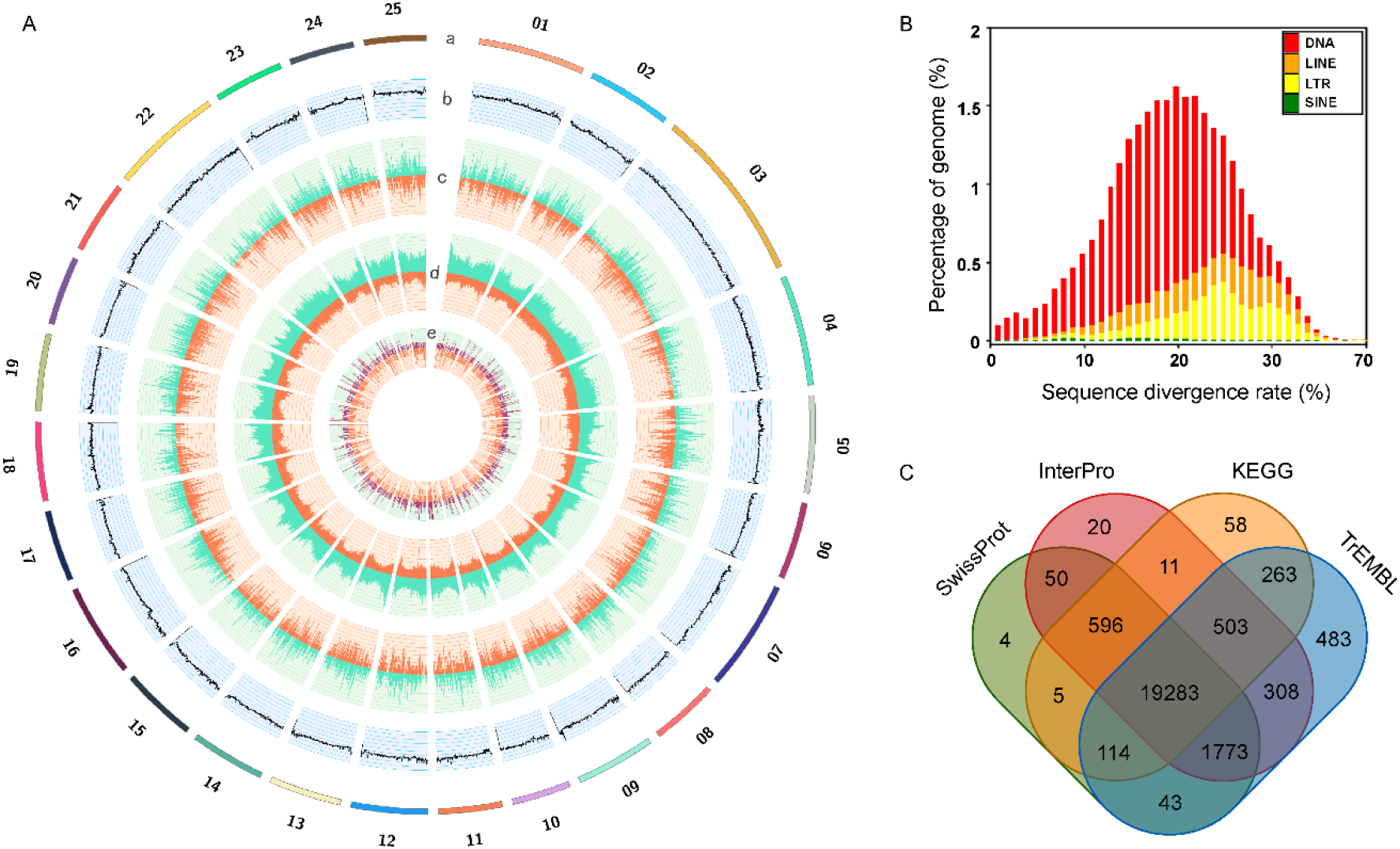
Genome Features of *P. huangchuchieni*. A, circos plot of the genome. From outer to inner circles: (a) Lines represent *P. huangchuchieni* chromosomes; (b) GC content; (c) Gene density; (d) Repeat element density; (e) ncRNA density. (b-e) are drawn in non-overlapping 100 kb sliding windows. B, Distribution of divergence rate for each type of TE in *P. huangchunchieni* genome. The divergence rate was calculated based on the identified TE elements from the Repbase. C, Venn diagram of the functionally annotated genes.

A previous study in *C. carpio* indicated that one of the subgenomes (B subgenome) may be derived from a diploid lineage in Barbinae, including genera *Poropuntius* and *Onychostoma* (Xu et al., 2019). The complete reference genome of *Onychostoma* (*O. macrolepis*) has been recently elucidated, thereby providing a comparable reference genome for the *P. huangchuchieni* genome. Both diploid genomes have the same karyotype (2n=50) and show high genome consistency and comparable genome size (Figure S1 and S2). The chromosome synthetic comparisons of *P. huangchuchieni* and the B subgenome in *C. carpio* also exhibited a high level of collinearity (Figure 3A), thus reflecting the high quality of *P. huangchuchieni* genome assembly as well as the progenitor-like character of *P. huangchuchieni* to *C. carpio*. The genome size of the B subgenome in *C. carpio* was significantly smaller than that of *P. huangchuchieni* (Figure S2), which might result from genome fusion and evolution in the allotetraploid genome.

**Figure 3.**
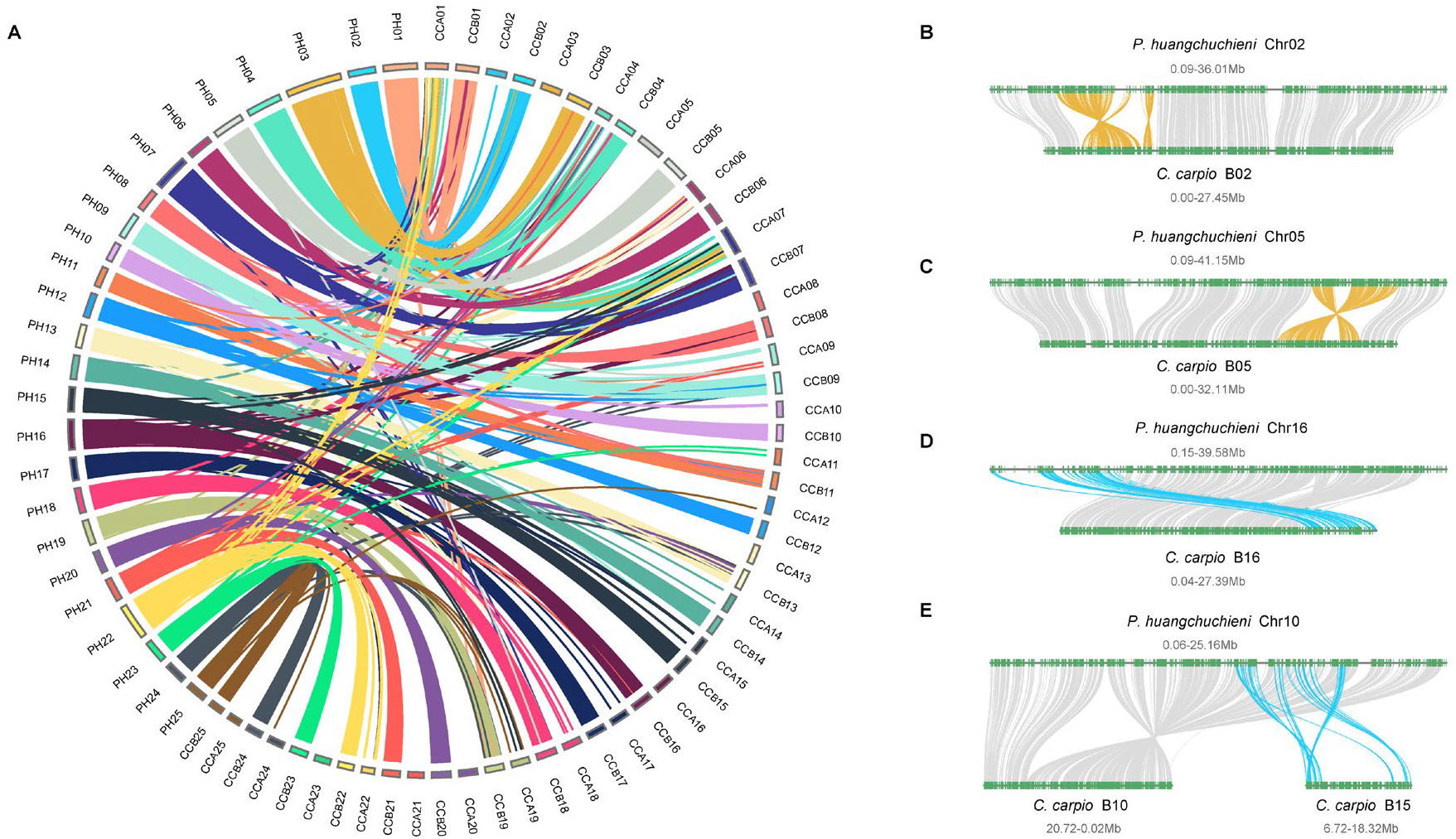
Synteny of *P. huangchuchieni* and *C. carpio*. A, *P. huangchuchieni* (PH) shows high consistence with the B subgenome of *C. carpio* (CC). B-E show structural variations of *P. huangchuchieni* and *C. carpio*. B and C show inversions, D and E show translocations and segmental duplications.

Structural variations were identified between *P. huangchuchieni* and the B subgenome in *C. carpio*, including inversions (Figure 3B and 3C), translocations (Figure 3D), and segmental duplications (Figure 3E), which provided insights into the chromosomal rearrangements that occurred following the WGD event in the allotetraploid *C. carpio* genome. Genomic rearrangements in plants have been broadly studied and it has been suggested that some of the rearrangements contribute to important agronomic traits, such as seed set and changes in flowering time (Osborn et al., 2003; Pires et al., 2004; Session et al., 2016; Yin et al., 2020). However, the underlying mechanisms for the rearrangements in polyploidy animals are poorly understood. With the closely related genome of *C. carpio*, the genome of *P. huangchuchieni* may be further used to identify the potential function of these regions and may promote a better understanding of the adaption and specific traits of *C. carpio*.

### Annotation of repetitive sequences

The consensus and non-redundant repetitive sets were obtained by a combination of known, novel and tandem repeats, which generated a total of 485.82 Mb of repetitive sequences, occupying more than 47% of the whole genome assembly (Figure 2B, Table 4, Table S8). The percentage of repetitive sequences in the *P. huangchuchieni* genome was similar to that of the other species in Barbinae, for example, *O. macrolepis* (409.96 Mb; 46.23%) (Sun et al., 2020), and was higher than that of *C. carpio* (36.42%) and *C. auratus* (39.49%) (Luo et al., 2020; Xu et al., 2019). DNA transposons are the most abundant repetitive elements, spanning at least 196.46 Mb or 19.25% of the genome, the number of which in the closely related *O. macrolepis* was 27.36% (Table 4, Table S9), intuitively larger than that of the *C. carpio* (Sun et al., 2020; Xu et al., 2019). The repetitive sequences also comprised of long interspersed elements in 49.10 Mb (LINEs; 4.81%), short interspersed nuclear elements in 2.46 Mb (SINEs; 0.24%) and long terminal repeats in 51.56 Mb (LTRs; 5.05%) (Table 4, Table S9). The differences between the diploid and the allotetraploid may be caused by hybridization and WGD, and transposable elements may be responsible for the genome size divergence (Marburger et al., 2018; Ramachandran, 2018; Talla et al., 2017).

**Table 4.** Statistic of transposable element in the genome.

| Type | Rebase TEs |  | TE protiens |  | *de novo* |  | Combined TEs |  |
| --- | --- | --- | --- | --- | --- | --- | --- | --- |
|  | Length<br>(Bp) | % in<br>genome | Length<br>(Bp) | % in<br>genome | Length<br>(Bp) | % in<br>genome | Length<br>(Bp) | % in<br>genome |
| **DNA** | 187,072,330 | 18.33 | 18,903,016 | 1.85 | 30,351,360 | 2.97 | 196,462,783 | 19.25 |
| **LINE** | 39,956,585 | 3.92 | 30,295,503 | 2.97 | 30,948,966 | 3.03 | 49,100,608 | 4.81 |
| **SINE** | 2,460,536 | 0.24 | 0 | 0 | 0 | 0 | 2,460,536 | 0.24 |
| **LTR** | 41,553,946 | 4.07 | 26,076,846 | 2.56 | 27,100,222 | 2.66 | 51,555,105 | 5.05 |
| **Other** | 7,335 | 0 | 0 | 0 | 0 | 0 | 7,335 | 0 |
| **Unknown** | 0 | 0 | 0 | 0 | 349,802,298 | 34.28 | 349,802,298 | 34.28 |
| **Total** | 264,770,429 | 25.95 | 75,124,080 | 7.36 | 428,915,511 | 42.04 | 485,818,804 | 47.61 |

### Prediction of protein-coding genes

Integrating *de novo*, homology searching and transcriptome-assisted predictions (transcriptomic data of 12 tissues listed in Table S10), we successfully generated a non-redundant gene set comprising of 24,099 protein-coding genes (Table S11). The statistics of the predicted gene models were compared with other teleost species, such as *D. rerio* and *O. macrolepis*, which showed similar distribution patterns in mRNA length, CDS length, exon length, intron length and exon number (Figure S3, Table S12).

Functional annotation was performed by comparing the protein sequences in several public gene databases. As a result, 90.74%, 86.45% and 94.49% of the predicted genes had positive hits in the SwissProt, KEGG, and TrEMBL databases, respectively. We also identified protein domains in multiple databases, and 93.55% and 66.55% of the predicted genes were annotated in InterPro and GO database, respectively. Finally, 23,514 genes (97.57% of all predicted genes) were successfully functional annotated in at least one of these databases (Figure 2C, Table S11, Figure S4).

Additionally, 378 miRNAs, 2472 tRNAs, 2,702 rRNAs and 814 snRNAs were identified by *de novo* or homology-based annotation (Table S13). The content of non-coding RNAs varied considerably, which was also observed between the closely related Barbinae (Sun et al., 2020). The underlying basis of these differences should be further investigated through multiple approaches. The genome features of *P. huangchuchieni* are illustrated in the circos plot (Figure 2A). The innermost circle shows the ncRNA distribution on different chromosomes (Figure 2A-e).

### Analysis of phylogenetic relationships and estimation of divergence time

To reveal the phylogenetic relationships among *P. huangchuchieni* and other species, we identified 992 single-copy ortholog families from nine representative species and aligned the protein sequences to generate a phylogenetic tree. The results showed that *P. huangchuchieni* clustered within the family Cyprinidae, together with *O. macrolepis, C. carpio, C. auratus* and *S. graham*, which was consistent with the phylogenetic relationship reported previously (Wang, Gan, Li, Chen, & He, 2016; Xu et al., 2019).

As for the divergence-time estimation, *P. huangchuchieni* was found to be separated from the B subgenomes of *C. carpio, C. auratus* and *S. grahami* approximately 12.25 Mya, which was in accordance with previous estimates of the allotetraploid genome origin and speciation time of the *C. carpio* genome using TE divergence (Xu et al., 2019). The divergence time of *P. huangchuchieni* and A subgenome was estimated at around 17.34 Mya (Figure 4). The time scale estimated was more likely to be in the Pliocene epoch, suggesting a relationship between polyploidy origin and dramatic climate changes during the QTP uplifting.

**Figure 4.**
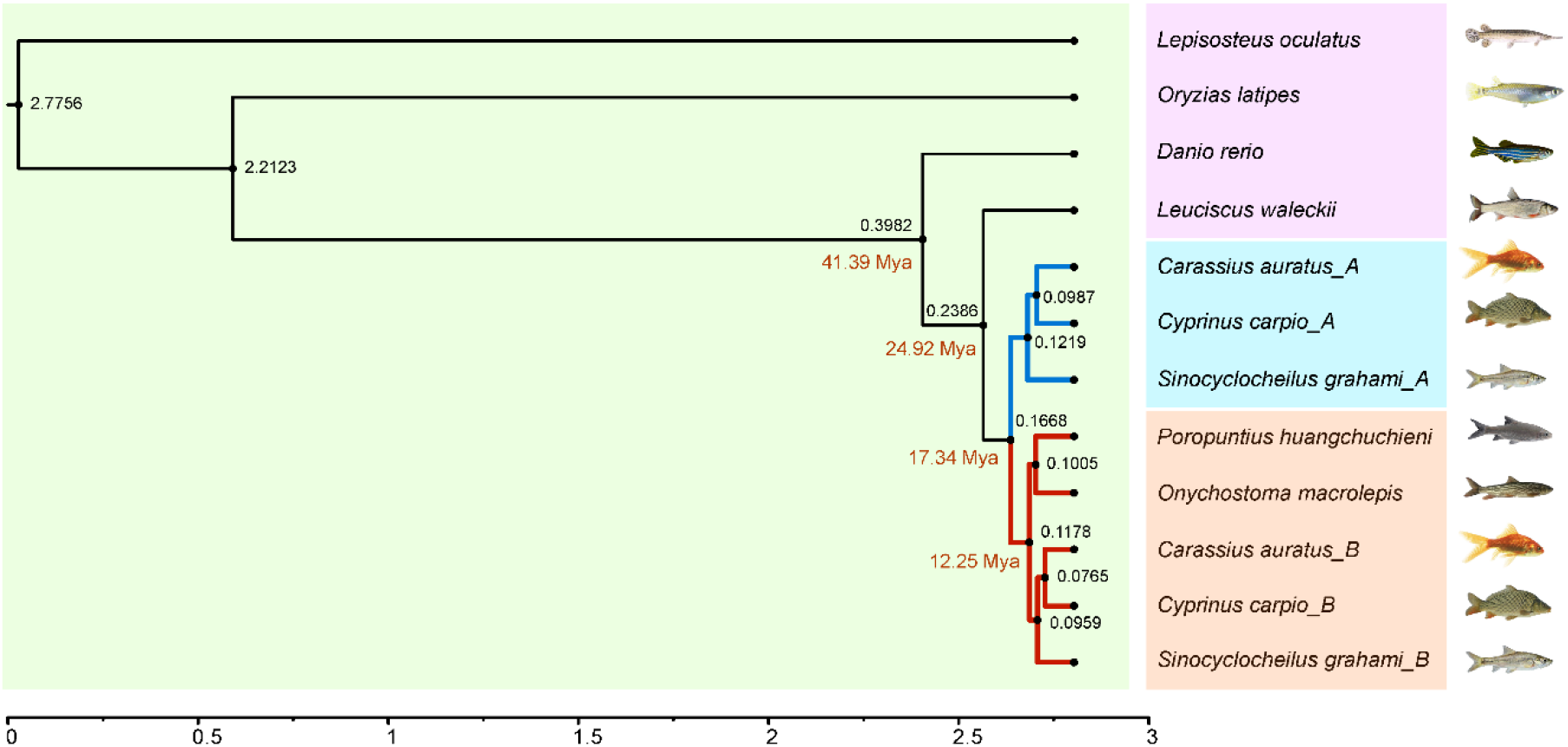
Phylogenetic tree of nine vertebrate genomes constructed using 992 single copy orthologous genes. Orthologous genes of the allotetraploid species were retrieved from the divided subgenomes of the species based on the genome similarity to *P. huangchuchieni*.

### Homoeologous gene triplets between *P. huangchuchieni* and *C. carpio*

Previously, we compared the gene contents of the allotetraploid of *C. carpio* and the diploid genome of *C. idella*, and finally developed 8,291 orthologous gene triplets that retained one copy in the diploid genome and one copy in each of the two homoeologous chromosomes in the two subgenomes of the *C. carpio* genome (Xu et al., 2019). For estimation of homoeologous gene triplets and gene loss, identification of nearly all genes in the analyzed genome is crucial, and detection in the closely related species is ideal. The high-quality genomes of both *P. huangchuchieni* and *C. carpio* provide the tool to develop a more thorough homoeologous gene set for further evolutionary studies.

In the present study, with information on the closest relationship between the diploid progenitor *P. huangchuchieni* and the allotetraploid *C. carpio*, we identified a total of 17,474 orthologous genes through the RBH approach. This included 13,389 homoeologous gene pairs in both the two homoeologous chromosomes in *C. carpio* with only one orthologous gene in the parental chromosomes in *P. huangchuchieni* (Table 5), which was 1.5 times than that reported for the previous gene sets. These homoeologous gene pairs retained in duplicate were not distributed evenly among Gene Ontology and were enriched in essential functional categories, such as RNA metabolic process (GO: 006396), cell growth (GO:0016049), lipid biosynthetic process (GO:0008610) (Figure 5). This is consistent, to some extent, with the previous identified 8,291 gene sets (Xu et al., 2019), thereby indicating that these genes were essential for the tetraploid *C. carpio*, and therefore a loss from the tetraploid genome was not possible (Albalat & Cañestro, 2016). On the other hand, the surviving duplicated genes might have occurred functional diversification, which was thought to be a major feature of the long-term evolution of polyploids (Blanc & Wolfe, 2004). As for the single-copy genes in *C. carpio*, the number was slightly larger than that reported previously, including 1,411 single-copy genes only in the B subgenome and 939 only in the A subgenome in *C. carpio.* A large fraction of the single-copy genes in both subgenomes belongs to functional GO categories such as DNA and RNA metabolic processes (GO:0006259, GO:0016070), DNA repair (GO:0006281), and response to stress (GO:0006950), which have been previously observed in yeast and plants (Aravind, Watanabe, Lipman, & Koonin, 2000), indicating that these genes are dispensable to some extent, and are therefore susceptible to loss because the loss has no or only a slightly negative impact on fitness (Albalat & Cañestro, 2016). An alternative hypothesis is that single-copy paralogs are retained towards genes evolve slowly or reciprocal gene loss is more likely to occur between duplicated genes with indistinguishable functions (Scannell, Byrne, Gordon, Wong, & Wolfe, 2006). Interestingly, single-copy genes retained only in B subgenome were likely to be related to GO terms such as RNA and ribosomal methylation (GO:0001510, GO:0032259), which was not observed when *C. idella* was used as the diploid reference genome (Figure 5). Our comparison of the ka, ks and ka/ks divergence between orthologs of *P. huangchuchieni* and *C. carpio* revealed that single-copy genes in *C. carpio* had much higher ka/ks values than the homoeologous gene pairs (Figure 5C), suggesting that the singleton genes experienced more relaxed purifying selection, and might accumulate more deleterious mutations, or their fast evolution resulted in the loss of one of the copies. The same phenomenon has been reported in other eukaryotic genomes (Davis & Petrov, 2004). Additionally, we found 442 and 672 homoeologous gene pairs located within the same subgenome in either A or B subgenome, which might be the result of homoeologous exchanges occurring post WGD event. These genes were intuitively related to categories including coenzyme metabolic processes (GO:0006732) and ion transport (GO:0006820). The full homoeologous gene set identified using the mostly related diploid progenitor *P. huangchuchieni* provides a broad view of the gene component in the allotetraploid genome, and is critical for future genome evolution and adaptation analysis in the allotetraploid *C. carpio*.

**Table 5.** Homoeologous genes between *C. carpio* and the reference diploid species.

| Category | PH_ortholog | CID_ortholog |
| --- | --- | --- |
| Ref:CCA:CCB=1:0:1 | 1,411 | 1,220 |
| Ref:CCA:CCB=1:1:0 | 939 | 915 |
| Ref:CCA:CCB=1:0:2 total | 672 | 110 |
| Ref:CCA:CCB=1:0:2 oneChr | 235 | 88 |
| Ref:CCA:CCB=1:0:2 twoChr | 437 | 22 |
| Ref:CCA:CCB=1:2:0 total | 442 | 126 |
| Ref:CCA:CCB=1:2:0 oneChr | 187 | 102 |
| Ref:CCA:CCB=1:2:0 twoChr | 255 | 24 |
| Ref:CCA:CCB=1:1:1 total | 14,010 | 8,353 |
| Ref:CCA:CCB=1:1:1 paired | 13,389 | 8,291 |
| Ref:CCA:CCB=1:1:1 non-paired | 543 | 62 |

The second and third column were the gene number using *P. huangchuchieni* and *C. idella* as the diploid reference.

**Figure 5.**
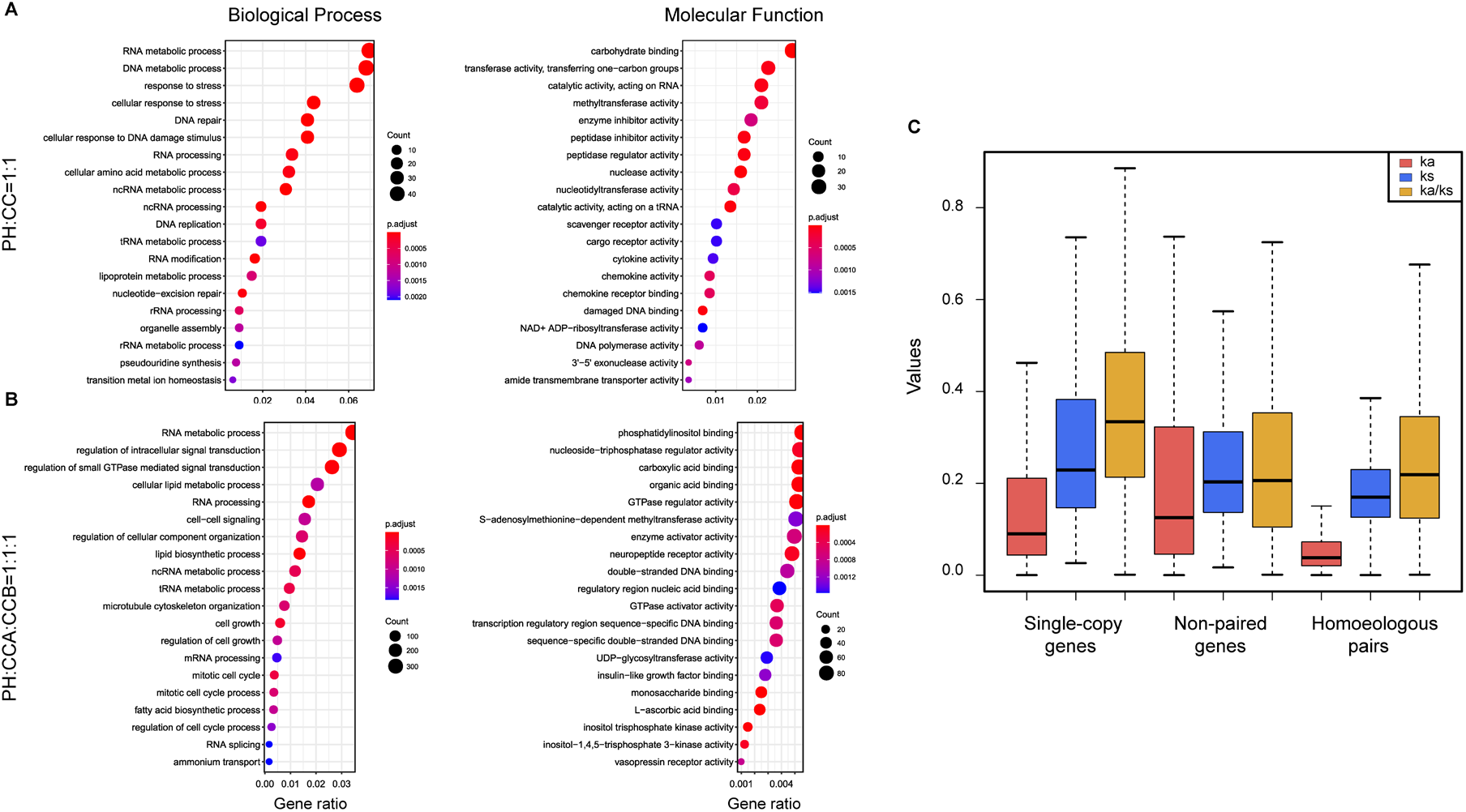
Significant GO terms of homoeologous genes in *C. carpio* and the selection pressure of orthologs between *P. huangchuchieni* and *C. carpio*. A, Top twenty GO terms of the single-copy genes in *C. carpio;* B, Top twenty GO terms of the Homoeologous gene pairs in *C. carpio*. C, Boxplots shows ka, ks, and ka/ks values for duplicable and singleton gene sets in *C. carpio*.

### Conclusion

In the present study, we provide a chromosomal-scale genome assembly and annotation of the diploid progenitor-like species of *C. carpio* by the integration of Nanopore long-read sequencing, Hi-C technology and Illumina short reads. Phylogenetic and syntenic analyses were performed to assess the evolutionary relationships of representative species. Orthologous genes between *P. huangchuchieni* and *C. carpio* were thoroughly analyzed, thus providing a broad view of the gene component in the allotetraploid genome after WGD. The high-quality genome data of *P. huangchuchieni* will provide a genomic and evolutionary connection between diploid and allotetraploid cyprinid species and thereby promote an understanding of the evolution and adaptation of cyprinid fish species.

## MATERIALS AND METHODS

### Ethics statement

Animal treatments in this study were conducted following the regulations of the Guide for Care and Use of Laboratory Animals and approved by the Committee of Laboratory Animal Experimentation at College of Ocean and Earth Sciences, Xiamen University.

### Sample collection and genome size estimation

*P. huangchuchieni* used in this study were obtained from Yunnan Province and reared at the aquarium of College of the Ocean and Earth Sciences, Xiamen University. Muscle tissues from an adult female *P. huangchuchieni* (Figure 1) were immediately frozen in liquid nitrogen, then stored at −80°C until samples were prepared for High Molecular Weight (HMW) DNA extraction. HMW DNA was extracted using a QIAGEN DNeasy Blood & Tissue Kit (QIAGEN, Shanghai, China). DNA quality was assessed and quantified using 1% agarose gel, NanoDrop™ One UV-Vis spectrophotometer, Qubit^®^ 3.0 Fluorometer and the Agilent 2100 Bioanalyzer. Paired-end sequencing libraries were constructed using the Illumina TruSeq Nano DNA Library Prep Kit, and sequenced on the Illumina HiSeq X Ten platform to generate sequences with a 150-bp read length. Quality control of raw sequencing reads was performed using Trimmomatic program v0.33 to remove adaptors and low-quality reads. Clean reads were applied to estimate the genome size and heterozygosity based on the 17-mer frequency distribution.

### Oxford Nanopore sequencing, assembly and correction

HMW DNA extracted from the same individual was used for long-read Nanopore sequencing. DNA fragments were size-selected with a Blue Pippin System. A Nanopore library was prepared using the Ligation Sequencing Kit (SQK-LSK109, ONT, USA) following the manufacturer’s instructions, and sequenced on flow cells of a PromethION sequencer. Base-calling was performed onboard the PromethION instrument using MinKnow versions 2.0-2.2 (ONT, USA). Nanopore clean reads were initially corrected using Canu, and then de novo assembled with Smartdenovo (https://github.com/ruanjue/smartdenovo). To further improve the accuracy of the assembly, two rounds of self-correction and two rounds of consensus correction were performed using ONT clean reads and Illumina short reads with Racon and Pilon.

### Chromosome assembly by Hi-C

To further generate a chromosomal-level assembly of the *P. huangchuchieni* genome, a high-throughput chromosome conformation capture technology (Hi-C) library was constructed using samples from the same fish. Hi-C libraries were sequenced on the Illumina HiSeq X Ten platform using a PE-150 module. Quality control of Hi-C raw data was performed using HiC-Pro. Hi-C clean reads of *P. huangchuchieni* were mapped to the Nanopore reference genome using Juicer to compare the Hi-C reads to the draft assembled sequence. Following this, low-quality reads were filtered out to build inter/intra-chromosomal contact maps. The Hi-C interactions were used as information for contig proximity. We then used the 3D-DNA pipeline to scaffold the *P. huangchuchieni* genome to 25 chromosomes.

### Repeat sequences annotation

Repeat sequences in the genome mainly consist of tandem and interspersed repeats. The latter is also known as a transposable element, including DNA transposons and retrotransposons. Two strategies were used to detect repeat sequences in the *P. huangchuchieni* genome. First, a homology-based detection was conducted using RepeatMasker and RepeatProteinMask (Chen, 2004) based on sequence similarity to known repeat sequences in the Repbase database (v.21.11, http://www.girinst.org/repbase). Second, a *de novo* repeat library was constructed by Piler (Edgar & Myers, 2005), RepeatScout (Price, Jones, & Pevzner, 2005), RepeatModeler (Chen, 2004) (http://www.repeatmasker.org/RepeatModeler/), and *de novo* annotation was performed using through Repeatmasker, Tandem Repeats Finder v4.09 (Benson, 1999) (http://tandem.bu.edu/trf/trf.html) and LTR_FINDER v1.06 (Xu & Wang, 2007) (http://tlife.fudan.edu.cn/ltr_finder/).

### The prediction and functional annotation of protein-coding genes

Protein-coding genes were predicted through a combination of a homology-based approach, *de novo* prediction and transcriptome-based approach. Protein sequences from *Danio rerio, Megalobrama amblycephala* and other representative species were aligned to the *P. huangchuchieni* genome using TblastN (E-value <= 1e-5). The BLAST hits were conjoined by Exonerate v2.2.0 (Slater & Birney, 2005) and Genewise v2.4.1 (Birney, Clamp, & Durbin, 2004) for best sequence alignments. For *de novo* prediction, Augustus (Stanke et al., 2006), GlimmerHMM (Majoros, Pertea, & Salzberg, 2004), SNAP (Johnson et al., 2008), Genscan (Burge & Karlin, 1997), FgeneSH (Solovyev, Kosarev, Seledsov, & Vorobyev, 2006), and GeneID v1.4.4 (Blanco, Parra, & Guigó, 2007) were used to predict the genes in the repeat-masked genome sequences. For transcriptome-based prediction, RNA-seq reads from intestines, liver, muscle, brain, spleen, skin, gill, kidney, head-kidney, blood, gonad and heart were aligned to the *P. huangchuchieni* genome using HISAT2 and Tophat2 (Kim et al., 2013). Transcripts were then assembled using Cufflinks and StringTie (Trapnell et al., 2010). Finally, gene predictions from the *homology-based, de novo*, and RNA-Seq-based evidence were merged to integrate a non-redundant and comprehensive consensus gene set using GLEAN (https://sourceforge.net/projects/glean-gene/), EVM (Haas et al., 2008), Maker and an in-house Ensembl-like pipeline.

To achieve the functional annotation, the predicted protein sequences were aligned against the public databases, including SwissProt (http://www.uniprot.org/), TrEMBL (http://www.uniprot.org/), KEGG (Kyoto Encyclopedia of Genes and Genomes, http://www.genome.jp/kegg/) and BLASTP (E-value<=1e-5). Additionally, protein motifs and domains were annotated by searching the InterPro (https://www.ebi.ac.uk/interpro/) and Gene Ontology (GO) databases using InterProScan (v4.8) (Zdobnov & Apweiler, 2001).

Benchmarking Universal Single-Copy Orthologs (BUSCO, RRID:SCR_015008) were used to assess the genome completeness and the accuracy of gene prediction by searching the predicted gene sets for the single-copy genes conserved in Actinopterygii.

### Non-coding RNA annotation

Non-coding RNAs include rRNAs, tRNAs, snRNAs, and miRNAs. The tRNAs in *P. huangchuchieni* genome were predicted based on the structural characteristics by tRNAscan-SE v1.3.1. The rRNAs were highly conserved among species and could be predicted by alignment against closely related species through BLASTN. INFERNAL included in Rfam (http://infernal.janelia.org/) was used for screening miRNAs and snRNAs in the genome.

### Phylogenetic tree construction and divergence time estimation

The phylogenetic relationships between *P. huangchuchieni* and other representative species were constructed using orthologs from single-copy gene families. The dataset included 992 protein sequences with one copy in the diploid *P. huangchuchieni, Onychostoma macrolepis, Danio rerio, Oryzias latipes, Leuciscus waleckii, Lepisosteus oculatus*, and two copies in each of the subgenomes of the tetraploid *C. carpio, Carassius auratus, Sinocyclocheilus grahami, Lepisosteus oculatus*. Protein sequences were aligned using BLASTP and phylogenetic analyses were conducted using RAxML v 8.2.12. The MCMCTREE software in the PAML package was used to estimate the divergence time of *P. huangchuchieni* from other species.

### Syntenic analysis between *P. huangchuchieni* and the closely related cyprinid species

To show global collinearity, Mummer v3.23 was adopted to conduct genome-wide sequence comparisons between the diploid *P. huangchuchieni* and *O. macrolepis* or the allotetraploid *C. carpio* genome sequences. Coordinates are displayed as circos plot. Detailed collinearity between *P. huangchuchieni* and the *C. carpio* B subgenome was provided by LASTZ v1.03.54. Furthermore, orthologous gene pairs between *P. huangchuchieni* and the *C. carpio* B subgenome and syntenic gene pairs were identified using the MCscan toolkit implemented in python [https://github.com/tanghaibao/jcvi/wiki/MCscan-(Python-version)] with a minimum block size of four genes. Macro- and microsynteny dot plots of structural variation (SV) were generated in Python using scripts from MCscan.

### Identification ofthe conserved homoeologous gene pairs and kaks analysis

The fully annotated protein-coding genes of *P. huangchuchieni* were used as a diploid reference to identify the conserved homoeologous gene pairs in the tetraploid genome of *C. carpio*. Ortholog prediction for genome-scale datasets was performed using a reciprocal-best-BLAST-hits (RBH) approach based on the all-against-all similarities using BLASTP. Orthologous triplets with a 1:1:1 relationship in the *P. huangchuchieni* genome and the A and B subgenomes of *C. carpio* were identified from two genomes. The orthologous gene pairs in the *C. carpio* genome were then defined as homoeologous genes that were derived from the latest whole genome duplication (WGD) event. Gene loss was observed for genes with one copy in *P. huangchuchieni* genome and only one orthologous gene in either of the two subgenomes.

The nonsynonymous (Ka) and synonymous (Ks) substitution rates of each orthologous gene pair between *P. huangchuchieni* and *C. carpio* were calculated using ParaAT (Parallel Alignment and back-Translation) and Muscle software were used to prepare the input files for KaKs_Calculator2.0 (Zhang et al., 2012).

### Data Accessibility

All genomic sequence datasets of *P. huangchuchieni* can be found on NCBI under Bioproject number PRJNA511031 (https://dataview.ncbi.nlm.nih.gov/object/PRJNA511031), the National Genomics Data Center under Bioproject number PRJCA002855 (https://bigd.big.ac.cn/gsub/submit/bioproject/subPRO004206/overview), Dryad (https://datadryad.org/stash/dataset/doi:10.5061/dryad.crjdfn32p) and Figshare (https://figshare.com/articles/dataset/Chromosome-level_genome_assembly_of_Poropuntius_huangchuchieni/12793595).

## Conflicts of interests

The authors declare no competing interests.

## Acknowledgements

This work was supported by the National Key Research and Development Program of China (2019YFE012050), the National Natural Science Foundation of China (grants 31872561), the Fundamental Research Funds for the Central Universities, Xiamen University (grants 20720180123 & 20720160110). We thank BGI Shenzhen for complimentary genome sequence collection on Nanopore PromethION platform under the “New Platform Cooperation” program in BGI with the help of Ms. Bin Geng.

## Author contributions

P.X conceived the project. L.C collected the sequencing samples, and extracted the DNA/RNA. L.C, B.C, C.L and Z.Z analyzed the data. W.Y collected the photograph of the fish. L.C wrote the manuscript. P.X, B.L and T.Z revised the manuscript. All authors reviewed and approved the final manuscript.

## Notes

### Competing Interest Statement

The authors have declared no competing interest.

