## Supplemental Information for "Chromosome-level genome of *Poropuntius huangchuchieni* provides a diploid progenitor-like reference genome for the allotetraploid *Cyprinus carpio*"

### Supplementary Figures

**Figure S1. Synteny of *P. huangchuchieni* and *O. macrolepis*. *P. huangchuchieni* (PH) showed high consistence with the *O. macrolepis* (OM).**

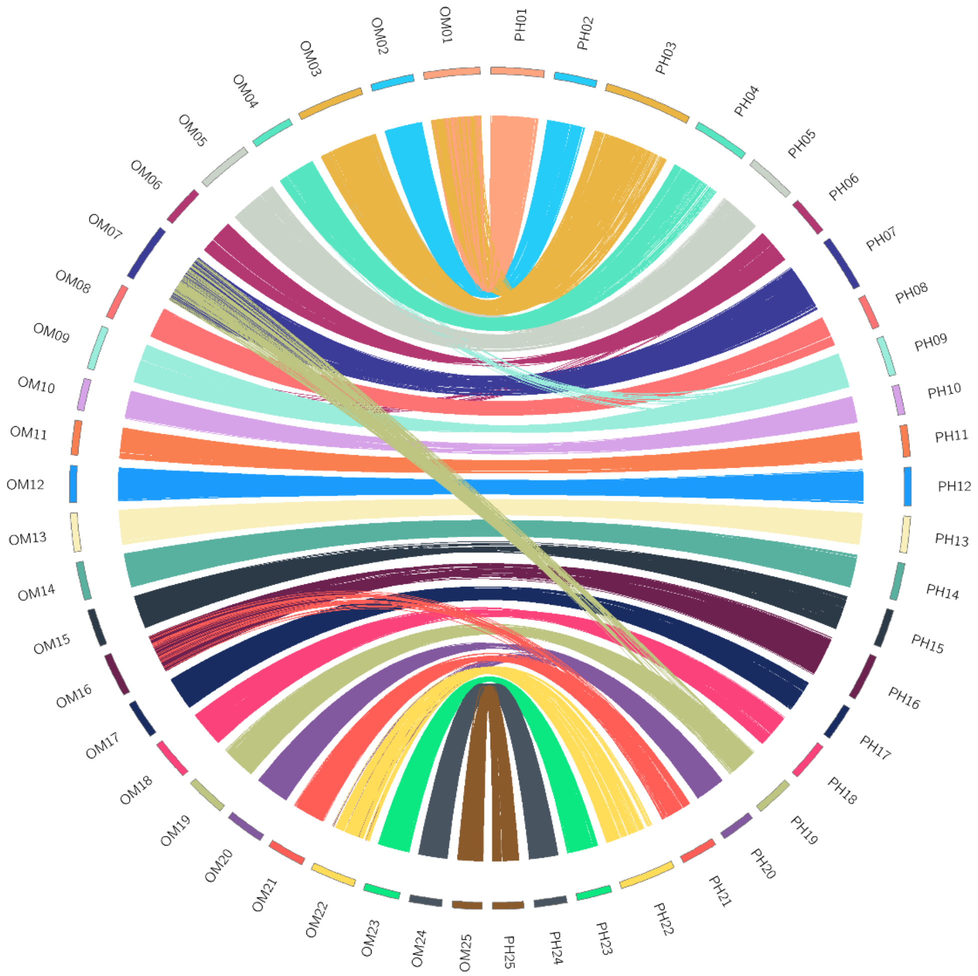

**Figure S2. Chromosome length comparison of *P. huangchuchieni*, *O. macrolepis* and *C. carpio*.**

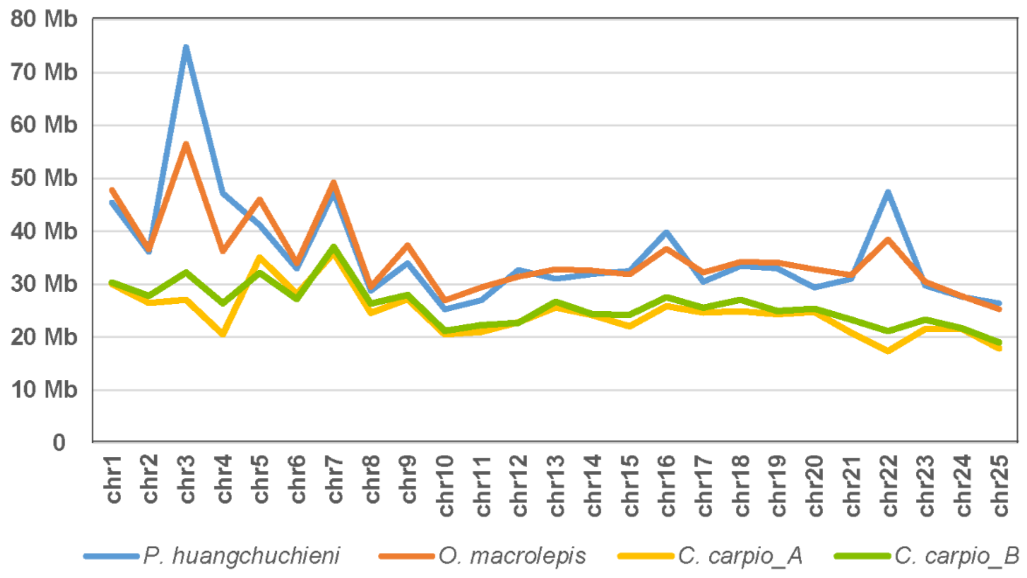

**Figure S3. The statistics of gene structure of *P. huangchunchieni* and the comparison with other teleost.**

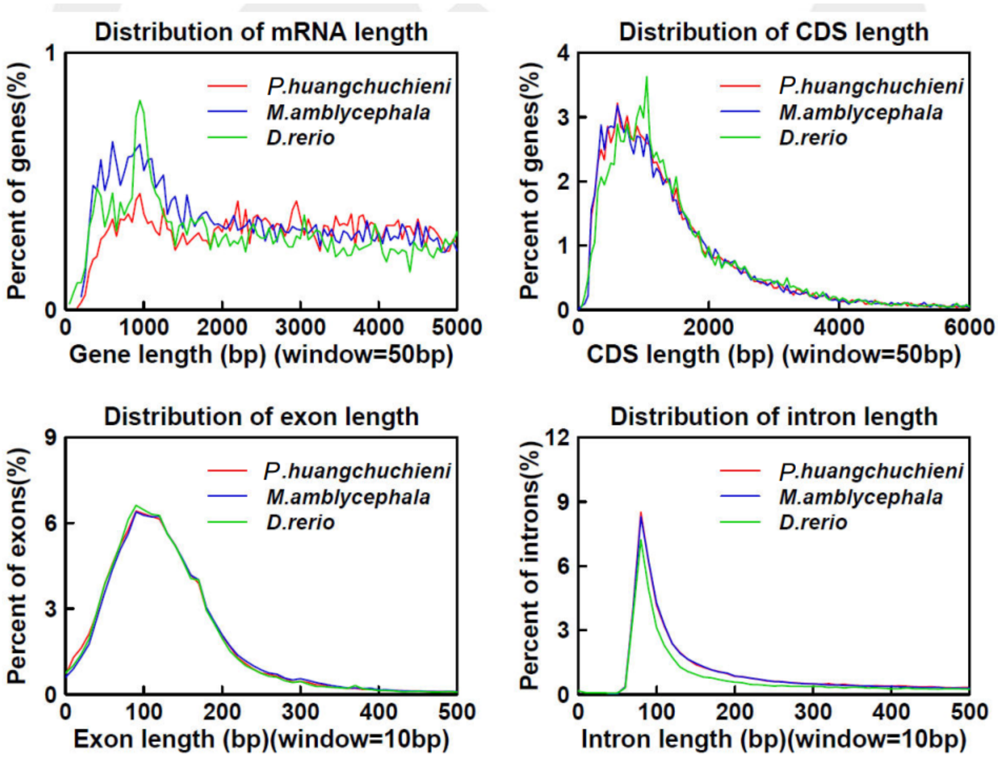

**Figure S4. GO and KEGG functional annotation of *P. huangchunchieni*.**

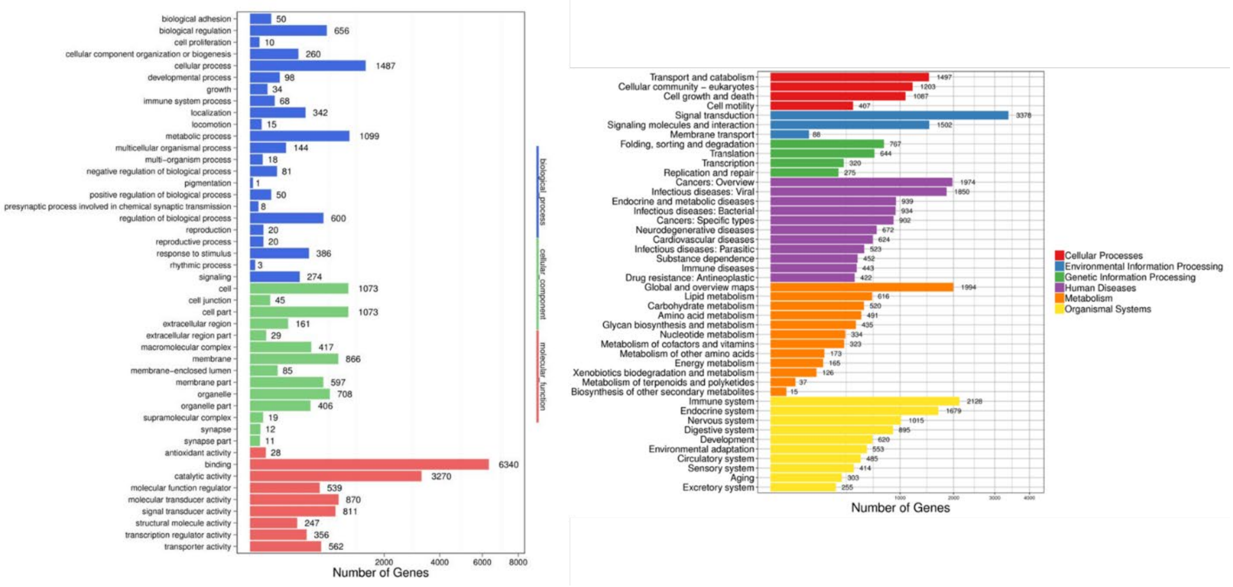

### Supplementary Tables

**Table S1. Statistics of illumina short-reads of *P. huangchunchieni* genomic survey.**

| Data | Reads | Bases(bp) | Read Length(bp) | Q30 | GC |
|---|---|---|---|---|---|
| Raw data | 382,339,662 | 57,350,949,300 | 150 | 89.77 | 37.41 |
| Clean data | 327,613,152 | 49,141,972,800 | 150 | 89.01 | 37.60 |

**Table S2. The data statistic of 17-mer analysis and Heterozygosity of *P. huangchunchieni* genome.**

| Sample | kmer | Depth | kmer number | kmer depth | Genome size(M) | Revised Genome size (M) | Heterozygosity (%) | Repeat rate (%) |
|---|---|---|---|---|---|---|---|---|
| Female | 17 | 47 | 43,900,162,368 | 47 | 934.05 | 911.37 | 0.68 | 48.84 |

**Table S3. The data statistic of Nanopore sequencing.**

| Sample | Total Reads | Total Base(GB) | Max Length(bp) | Mean Length(bp) | N50 Length(bp) | MeanQV |
|---|---|---|---|---|---|---|
| Raw data | 3,615,895 | 76.06 | 299,839 | 21,036 | 30,175 | 12.16 |
| Clean data | 3,162,952 | 74.78 | 299,805 | 23,641 | 30,598 | 12.17 |

**Table S4. Reads distribuation of Nanopore sequencing data.**

| Length | Reads Number | Total Length(bp) | Percent | Average Length(bp) |
|---|---|---|---|---|
| 5k~10k | 522,018 | 4,062,182,986 | 5.43% | 7,782 |
| 10k~20k | 1,212,973 | 17,359,315,510 | 23.22% | 14,311 |
| 20k~30k | 618,006 | 15,158,330,341 | 20.27% | 24,528 |
| 30k~40k | 350,936 | 12,125,660,579 | 16.22% | 34,552 |
| 40k~50k | 201,896 | 8,991,169,230 | 12.02% | 44,534 |
| 50k~60k | 114,774 | 6,257,611,074 | 8.37% | 54,521 |
| 60k~70k | 64,803 | 4,180,458,946 | 5.59% | 64,510 |
| 70k~80k | 36,118 | 2,691,845,981 | 3.60% | 74,529 |
| =>80k | 41,428 | 3,949,299,303 | 5.28% | 95,329 |

**Table S5. Summary of preliminary genome assembly using Nanopore sequencing data.**

| **Statistics type** | **Contig length(bp)** | **contig number** |
|---|---|---|
| **Max length** | 19,993,261 | |
| **N10** | 13,147,925 | 7 |
| **N20** | 9,500,847 | 17 |
| **N30** | 6,928,818 | 29 |
| **N40** | 5,103,178 | 47 |
| **N50** | 4,103,268 | 69 |
| **N60** | 2,354,835 | 103 |
| **N70** | 1,587,950 | 155 |
| **N80** | 951,368 | 238 |
| **N90** | 456,036 | 396 |
| **Total length** | 1,020,362,143 | |
| **number>=100bp** | 919 | 919 |
| **number>=2000bp** | 919 | 919 |

**Table S6. Statistics of Hi-C data.**

| Categories | Data size |
|---|---|
| Raw Paired-end Reads | 334,637,855 |
| Raw Bases(bp) | 100,391,356,500 |
| Clean Q30 Bases Rate (%) | 92.43 |
| Clean Paired-end Reads | 330,936,826 |
| Clean Bases(bp) | 99,281,047,800 |
| Clean Paired-end Reads Rate (%) | 98.89 |

**Table S7. Chromosome summary of *P. huangchuchieni*.**

| Chromosome | Scaffold Number | *P. huangchuchieni* |
|---|---|---|
| chr1 | 40 | 45,381,146 |
| chr2 | 31 | 36,035,969 |
| chr3 | 150 | 74,782,395 |
| chr4 | 82 | 47,136,578 |
| chr5 | 20 | 41,173,690 |
| chr6 | 22 | 32,930,180 |
| chr7 | 44 | 47,327,956 |
| chr8 | 13 | 28,731,048 |
| chr9 | 31 | 33,922,736 |
| chr10 | 21 | 25,216,553 |
| chr11 | 17 | 26,990,487 |
| chr12 | 42 | 32,666,103 |
| chr13 | 28 | 30,954,918 |
| chr14 | 19 | 31,888,916 |
| chr15 | 29 | 32,506,248 |
| chr16 | 48 | 39,809,036 |
| chr17 | 14 | 30,453,563 |
| chr18 | 39 | 33,408,662 |
| chr19 | 27 | 32,963,991 |
| chr20 | 16 | 29,339,261 |
| chr21 | 30 | 30,948,149 |
| chr22 | 120 | 47,409,594 |
| chr23 | 36 | 29,686,815 |
| chr24 | 21 | 27,609,575 |
| chr25 | 23 | 26,383,430 |
| Total | 963 | 895,656,999 |

**Table S8. Statistics of repetitive sequences in the P. huangchuchieni genome.**

| Type | Repeat Size (bp) | % of genome |
|---|---|---|
| Trf | 41,166,095 | 4.03 |
| Repeatmasker | 264,770,429 | 25.95 |
| Proteinmask | 75,124,080 | 7.36 |
| De novo | 439,923,381 | 43.11 |
| Total | 507,427,260 | 49.73 |

**Table S9. Comparison of transposable element in different genomes.**

| Type | *P. huangchuchieni* | | *O. macrolepis* | | *C. carpio* | | *C. auratus* | |
|---|---|---|---|---|---|---|---|---|
| | Length (Bp) | % in genome | Length (Bp) | % in genome | Length (Bp) | % in genome | Length (Bp) | % in genome |
| **DNA** | 196,462,783 | 19.25 | 242,634,604 | 27.36 | 199,140,970 | 13.97 | 326,579,818 | 21.19 |
| **LINE** | 49,100,608 | 4.81 | 43,172,542 | 4.87 | 105,515,036 | 7.40 | 88,268,888 | 5.73 |
| **LTR** | 51,555,105 | 5.05 | 38,664,341 | 4.36 | 96,119,903 | 6.74 | 89,602,232 | 5.81 |
| **SINE** | 2,460,536 | 0.24 | 2,804,360 | 0.32 | 1,673,649 | 0.27 | 9,493,708 | 0.62 |
| **Total** | 485,818,804 | 47.61 | 409,960,927 | 46.23 | 518,982,209 | 36.42 | 608,527,235 | 39.49 |

**Table S10. Statistics of data in transcriptome sequencing by Illumina platform.**

| Sample | Raw reads | Clean reads | Raw bases | Clean bases | Q30(%) | GC content(%) |
|---|---|---|---|---|---|---|
| Intestines | 55,236,326 | 54,529,562 | 8,285,448,900 | 8.18G | 94.85 | 47.94 |
| Liver | 49,010,228 | 47,541,104 | 7,351,534,200 | 7.13G | 95.66 | 48.39 |
| Muscle | 47,705,904 | 47,174,152 | 7,155,885,600 | 7.08G | 94.93 | 49.47 |
| Brain | 43,760,920 | 43,308,838 | 6,564,138,000 | 6.5G | 95.36 | 47.29 |
| Spleen | 48,599,034 | 48,027,448 | 7,347,112,800 | 7.2G | 91.99 | 48.43 |
| Skin | 49,923,296 | 49,412,964 | 7,488,494,400 | 7.41G | 94.78 | 48.38 |
| Gill | 51,725,152 | 50,977,286 | 7,758,772,800 | 7.65G | 95.4 | 47.05 |
| Kidney | 52,742,292 | 51,655,448 | 7,911,343,800 | 7.75G | 94.54 | 48.29 |
| Head kidney | 49,954,668 | 49,400,220 | 7,493,200,200 | 7.41G | 95.25 | 49.18 |
| Blood | 49,291,446 | 48,666,802 | 7,641,162,900 | 7.3G | 95.55 | 48.44 |
| Gonad | 45,476,130 | 44,923,624 | 6,821,419,500 | 6.74G | 95.34 | 47.63 |
| Heart | 54,396,420 | 53,758,508 | 8,159,463,000 | 8.06G | 94.83 | 47.84 |

**Table S11. Summary of gene functional annotation in different databases**

| **Values** | **Total** | **GO** | **Swissprot** | **KEGG** | **TrEMBL** | **Interpro** | **Overall** |
|---|---|---|---|---|---|---|---|
| **Number** | 24,099 | 16,037 | 21,868 | 20,833 | 22,770 | 22,544 | 23,514 |
| **Percentage** | **100%** | 66.55% | 90.74% | 86.45% | 94.49% | 93.55% | 97.57% |

**Table S12. Statistics of gene predictions in *P. huangchuchieni a*nd other species.**

| Species | Total number of gene | Average transcript length (bp) | Average CDS length (bp) | Average exons number per gene | Average exon length (bp) | Average intron length (bp) |
|---|---|---|---|---|---|---|
| *P. huangchunchieni* | 24,099 | 15,769.09 | 1,549.71 | 9.26 | 167.22 | 1,375.70 |
| *O.macrolepis* | 24,770 | 18,734.40 | 1,701.88 | 10.38 | 163.92 | 1,815.35 |
| *D. rerio* | 26,046 | 24,122.71 | 1,593.16 | 9.29 | 171.51 | 2,717.27 |
| *C. carpio* | 44,626 | 13 814.17 | 1,648.00 | 9.57 | 171.29 | 1244.38 |
| *S.grahami* | 42,109 | 14,530.83 | 1,625.70 | 9.24 | 175.89 | 1,565.69 |

Notes: *P. huangchunchieni*, *Poropuntius huangchuchieni*; *O.macrolepis*, *Onychostoma macrolepis*; *D. rerio*, *Danio rerio*; *C. carpio, Cyprinus carpio S. grahami*, *Sinocyclocheilus graham*.

**Table S13. Summary statistics of non-coding RNA.**

| Type | | Copy(w) | Average length(bp) | Total length(bp) | % of genome |
|---|---|---|---|---|---|
| **miRNA** | | 378 | 84.46 | 31,926 | 0.00313 |
| **tRNA** | | 2,472 | 76.86 | 189,993 | 0.01862 |
| **rRNA** | **rRNA** | 2,702 | 91.57 | 247,423 | 0.02425 |
| | **18S** | 32 | 184.91 | 5,917 | 0.00058 |
| | **28S** | 67 | 222.6 | 14,914 | 0.00146 |
| | **5.8S** | 2 | 156 | 312 | 0.00003 |
| | **5S** | 2,601 | 87 | 226,280 | 0.02218 |
| **snRNA** | **snRNA** | 814 | 149.7 | 121,853 | 0.01194 |
| | **CD-box** | 120 | 109.69 | 13,163 | 0.00129 |
| | **HACA-box** | 69 | 155.46 | 10,727 | 0.00105 |
| | **splicing** | 607 | 157.12 | 95,370 | 0.00935 |
